# Ephrin-A5 or EphA7 stimulation is anti-proliferative for human rhabdomyosarcoma *in vitro*

**DOI:** 10.1101/2024.12.23.629471

**Authors:** A. Cecchini, L. Ceccon, A. Chen, J. C. Schwesig, DDW Cornelison

## Abstract

Rhabdomyosarcoma (RMS) is a tumor which resembles skeletal muscle. Current treatments are limited to surgery and non-targeted chemotherapy, highlighting the need for alternative therapies. Differentiation therapy uses molecules that act to shift the tumor cells’ phenotype from proliferating to differentiated, which in the case of skeletal muscle includes exit from the cell cycle and potentially fusion into myofibers. We previously identified EphA7 expressed on terminally differentiated myocytes as a potent driver of skeletal muscle differentiation: stimulation of ephrin-A5-expressing myoblasts with EphA7 causes them to undergo rapid, collective differentiation. We therefore tested EphA7 as a candidate molecule for differentiation therapy on human RMS (hRMS) cell lines. Surprisingly, EphA7 had a lesser effect than ephrin-A5, a difference explained by the divergent suite of Ephs and ephrins expressed by hRMS. We show that in hRMS ephrin-A5 binds and signals to EphA8 and EphA7 binds and signals to ephrin-A2, and that Fc chimeras of both molecules are potent inhibitors of hRMS proliferation. These results identify key differences between hRMS and normal muscle cells and support further research into Eph:ephrin signaling as potential differentiation therapies.

**Summary statement:** This study identifies EphA7 and ephrin-A5 as external regulators of rhabdomyosarcoma proliferation, highlighting ephrin-A5 as a potential candidate for differentiation therapy in future cancer treatments.

## Introduction

Rhabdomyosarcoma (RMS) is a type of soft tissue cancer often arising during childhood that resembles skeletal muscle. Although 60% of patients diagnosed with RMS are effectively treated, the 5-year survival rate is only 33% in case of metastasis (Ognjanovic et al., 2009). RMS is divided in two main subtypes: embryonal (ERMS), which resembles embryonic muscle (Moore and Grossi, 1959), and alveolar (ARMS), which has morphology resembling pulmonary alveoli (Enterline and Horn, 1958). ERMS is characterized by multiple somatic mutations, with most significant ones being the loss of heterozygosity on chromosome 11 and deletions in the region containing at least three tumor suppressor genes (*IGF2*, *CDKN1C*, and *H19*) (Loh et al., 1992; Xia et al., 2002; Parham and Barr, 2013). ERMS represents ∼75% of RMS cases (Gurney et al., 1996) and is considered to be less aggressive compared to ARMS, with 5-year survival rates of 73.4% (Ognjanovic et al., 2009). ARMS tumors are characterized by translocation mutations that result in expression of fusions between the transcription activation domain of FOXO1 and the DNA-binding domains of PAX3 or (less frequently) PAX7, resulting in a tumor- promoting transcription factor (Galili et al., 1993; Shapiro et al., 1993; Barr, 2001). ARMS is more aggressive, with a 5-year survival rate of 47.8% (Ognjanovic et al., 2009).

RMS is categorized as a skeletal muscle tumor (Stout, 1946) and it is suggested to be the only tumor derived from a skeletal muscle lineage (Abraham et al., 2014), although RMS has also been described arising from another mesodermal lineage (adipose) (Hatley et al., 2012). RMS cells heterogeneously express satellite cell (muscle stem cells) markers such as Pax7 and Pax3 (Mansouri, 1998) as well as the muscle-specific transcription factors MyoD and myogenin (Sebire and Malone, 2003), although in normal muscle myogenin is exclusively expressed by terminally differentiated myocytes that have permanently exited the cell cycle.

Current treatment for RMS varies by tumor site and tumor type but is largely limited to conventional methods such as surgical removal, chemotherapy, or radiation. A more tailored approach to treatment would be differentiation therapy: this is particularly applicable in a muscle-like tumor because differentiated muscle is terminally postmitotic. The first molecule to be exploited for differentiation therapy was retinoic acid for the treatment of teratocarcinoma (Strickland and Mahdavi, 1978): it was observed that exposure to retinoic acid in the nM range caused morphological and metabolic changes in teratocarcinoma cells in *vitro*. Differentiation therapy via treatment with retinoids has been quite successful for hematopoietic tumors [reviewed in (Madan and Koeffler, 2021; Nagai and Ambinder, 2023)] and neuroblastoma [reviewed in (Makimoto et al., 2024)], and additional differentiation therapy molecules have been identified in the clinic, even for solid tumors [reviewed in (Cruz and Matushansky, 2012; Chen et al., 2020; Bar-Hai and Ishay-Ronen, 2022)].

The Eph receptor family is the largest RTK family in vertebrates. There are 14 members divided into two classes – EphA receptors (EphA1-8 and EphA10) and EphB receptors (EphB1-4 and EphB6) – which are categorized based on affinity for the two subclasses of membrane-bound ligands, ephrin-As and ephrin-Bs (Frisen et al., 1999). The ephrin-As (ephrin-A1-5) are tethered by a glycosylphosphatidlyinositol (GPI) anchor while ephrin-Bs (ephrin-B1-3) are transmembrane proteins (Davis et al., 1994; Pandey et al., 1995). Together, Ephs and ephrins constitute a bidirectional signaling system involved in contact-dependent cell-to-cell communication that can signal into the receptor (forward) and ligand (reverse) bearing cell or in both directions at the same time [reviewed in (Egea and Klein, 2007; Wu et al., 2019)].

Receptor-ligand binding is context-dependent and highly promiscuous: the EphAs bind to most or all of the ephrin-As, and the EphBs bind to most or all of the ephrin-Bs, and some interactions also occur across classes [reviewed in (Himanen et al., 2007; Dai et al., 2014)].

As Ephs are the largest family of RTKs in humans and are expressed in most tissues throughout development as well as being highly represented in stem cell lineages, it is unsurprising that dysregulated Eph:ephrin signaling has been shown to drive formation, progression, and/or metastasis of multiple tumor types [reviewed in (Chen, 2012; Kandouz, 2012; Pasquale, 2024)]. Comparatively little is known about possible activities of Ephs and ephrins in RMS: both EphBs and ephrin-Bs are upregulated in RMS (Berardi et al., 2008), EphA3 is expressed in a subset of

RMS cell lines and suppresses cell adhesion and migration (Clifford et al., 2008), and we recently noted that EphA1 is aberrantly localized to the nucleus of RMS in mouse, dog, and man (LaCombe et al., 2022).

We recently demonstrated that EphA7 presented on the surface of differentiated myocytes forces rapid collective differentiation of myoblasts via binding to ephrin-A5 on the myoblast cell surface, and also showed that an Fc chimera with the EphA7 extracellular domain is sufficient to induce myoblast differentiation *in vitro* (Arnold et al., 2020). We therefore asked if EphA7-Fc would also be sufficient to induce myogenic differentiation (and therefore halt proliferation) of RMS cells.

## Results

### EphA7-Fc and ephrin-A5-Fc treatment inhibits proliferation in hRMS cells

To ask if EphA7-Fc can inhibit proliferation in hRMS cells, we repeated the experiments that had previously demonstrated its efficacy in primary satellite cells on three hRMS cell lines [Rh41 and Ph30 (ARMS) and RD (ERMS)]; ephrin-A5 Fc was also used as a control.

Surprisingly, while EphA7-Fc inhibited proliferation in all three cell lines as measured by expression of Ki67, ephrin-A5 not only also inhibited proliferation but did so to a greater degree than EphA7-Fc (Figure 1).

**Figure 1.**
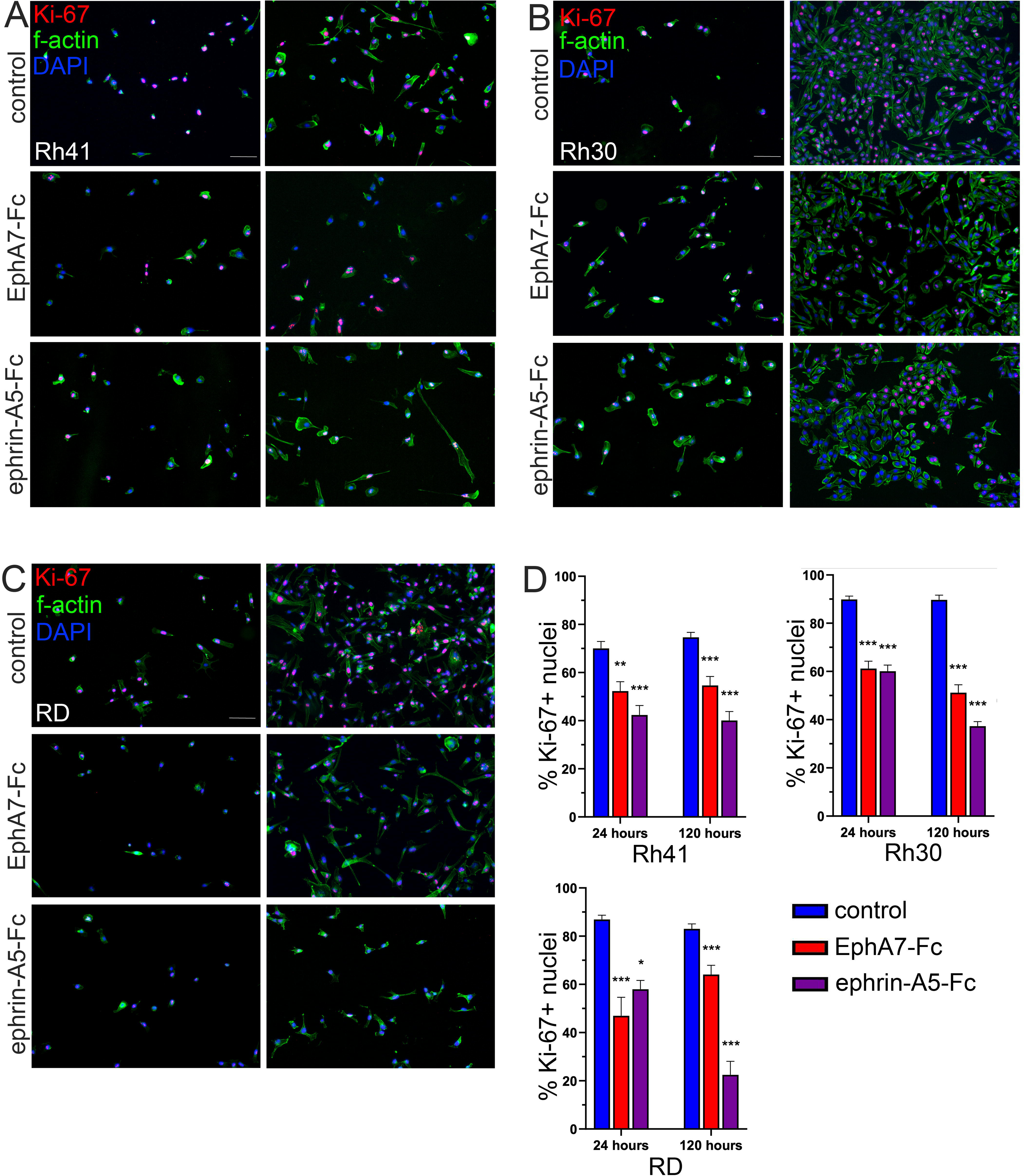
Treatment with EphA7-Fc and ephrin-A5-Fc decreases proliferation in hRMS cell lines. Rh41 ARMS (**A**), Rh30 ARMS (**B**), and RD (ERMS) (**C**) hRMS cell lines show decreased proliferation (Ki-67, red) (**D**) when cultured on coverslips functionalized with Fc chimeras of EphA7 ectodomain or ephrin-A5 compared to unprogrammed coverslips. Surprisingly, the effect of ephrin-A5-Fc is greater than that of EphA7-Fc in all cell lines, especially at 120 hours. Bars = 50 μm; *** = p<0.0001, ** = p<0.001, * = p<0.01.

### Expression and localization of EphA7 and ephrin-A5 in hRMS cells differs from *bona fide* myoblasts

Because of the unexpected results of ephrin-A5-Fc stimulation, we surveyed hRMS cells for expression of EphA7 and ephrin-A5 (which in skeletal muscle myoblasts during either development or regeneration are the ligand for ephrin-A5 and the receptor for EphA7, respectively) as well as the other mammalian Ephs and ephrins. We noted that while EphA7 is detectable in both ARMS cell lines, it is mislocalized to the *cis*-Golgi apparatus rather than the nucleus, and its expression in ERMS is greatly reduced (Figure 2 A-D). Moreover, expression of ephrin-A5 was not detectable in either ARMS or ERMS (Figure 2 A-C, E). These data suggested that EphA7-Fc could not be signaling via binding to ephrin-A5 in hRMS cells, and that the potential for ephrin-A5-Fc to be acting via binding to EphA7 was low, suggesting that they may be acting through other members of the family.

**Figure 2.**
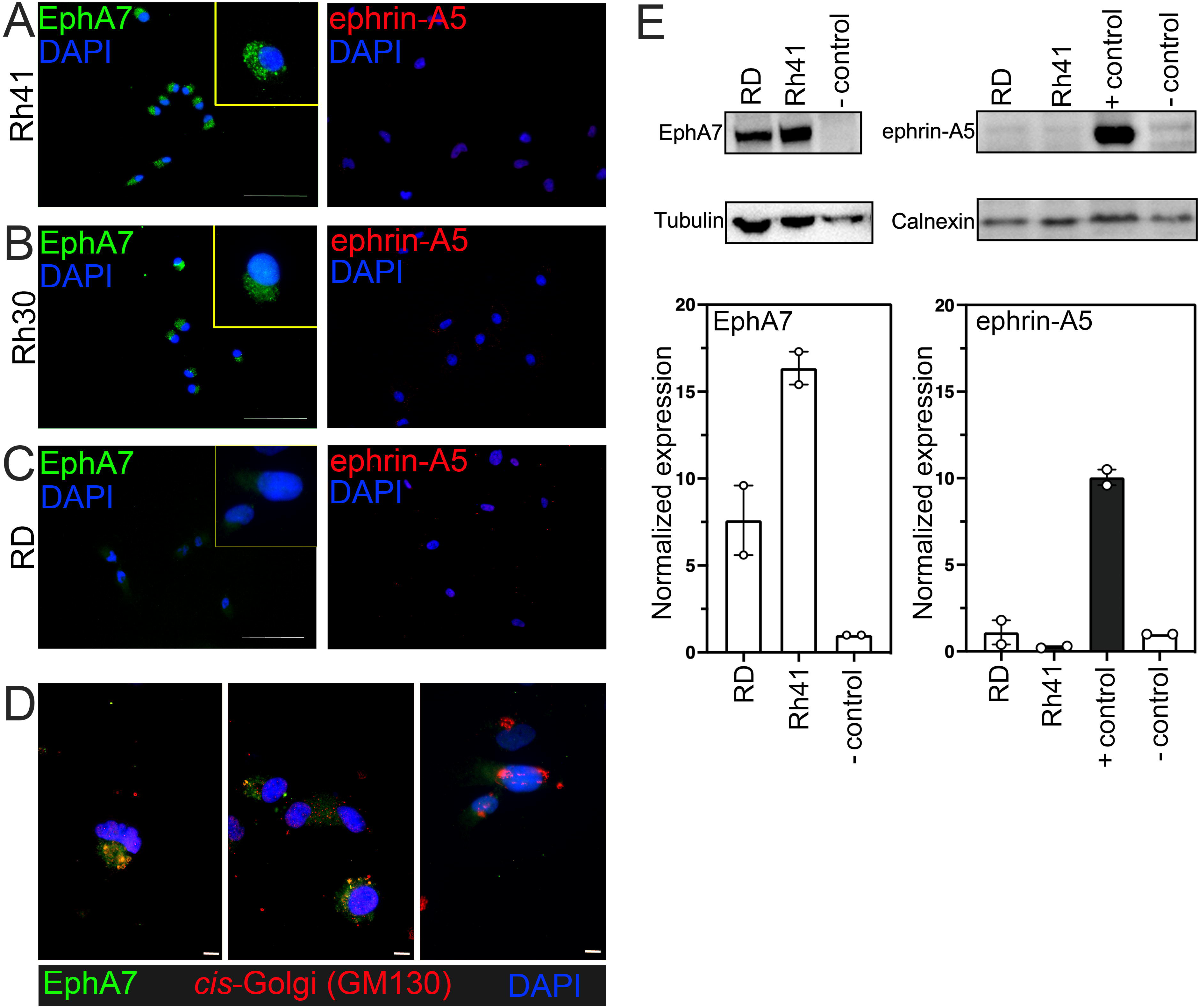
EphA7 and ephrin-A5 are either misexpressed or not expressed in hRMS. EphA7 protein (green) is not expressed at the plasma membrane of hRMS cells but in the perinuclear region in Rh41 (**A**) and Rh30 (**B**) and is also only weakly expressed in RD (**C**). Ephrin-A5 protein is not detectably expressed in any of the cell lines. Bars = 50 μm. EphA7 and GM130 (a *cis*-Golgi marker) colocalize in Rh41 cells, suggesting that EphA7 is aberrantly localized in the *cis*-Golgi apparatus in ARMS (**D**). Bars = 10 μm. Expression is confirmed in Western blot: EphA7 expression is decreased in RD cells compared to Rh41 cells by ∼5x, and ephrin-A5 was not detected (**E**); lysate from CRISPR-edited C2C12 cells which express neither ephrin-A5 nor EphA7 was included as a negative control and unedited C2C12 cell lysate was included as positive control.

An initial screening by RT-PCR (Supplemental Figure 1) showed detectable expression of ephrin-A2, ephrin-A3, ephrin-A5, and ephrin-B1 and EphA5, EphA6, EphA7, EphA8, EphB1, EphB2, EphB3, and EphB4 in at least one cell line. Based on these RT-PCR results, we performed immunocytochemistry for these Ephs and ephrins and noted expression of ephrin-A2, ephrin-A3, ephrin-B1, EphA6, EphA8, and EphB3 protein in all three cell lines (Figure 3). These members of the Eph:ephrin family are thus the primary candidates for interactions with either EphA7 or ephrin-A2.

**Figure 3.**
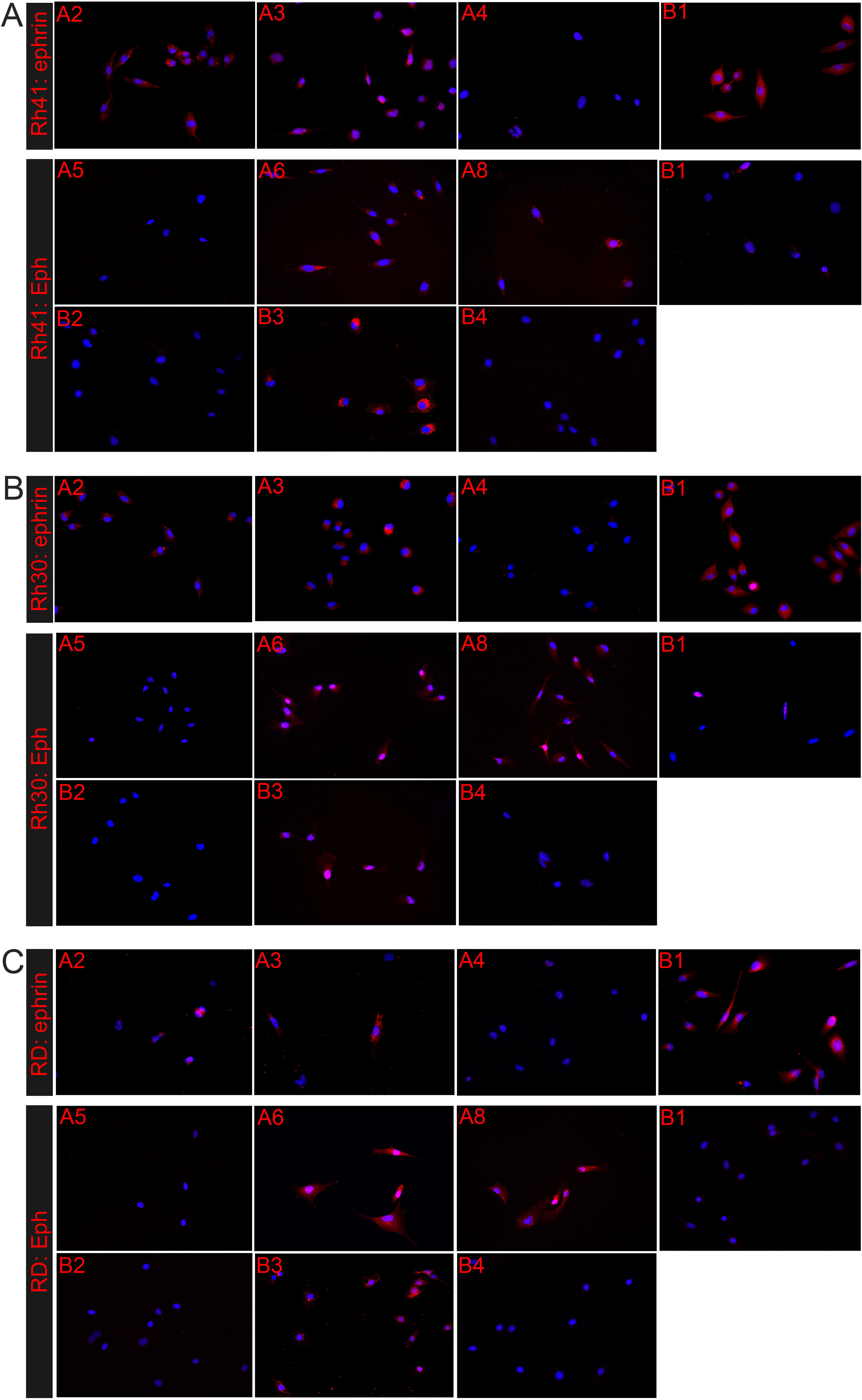
hRMS cell lines express a suite of Ephs and ephrins that is consistent among hRMS cell lines but differs from that of from myoblasts during development or regeneration. RT-PCR screen for all mammalian Ephs and ephrins shows that their expression in hRMS, while generally consistent across cell lines, differs from that of developing myoblasts or muscle satellite cell-derived myoblasts (**see data in Supplemental Figure 1**). Antibody staining for Ephs and ephrins detected by RT-PCR shows protein expression of ephrin-A2, ephrin-A3, and ephrin-B1 and of EphA6, EphA8, and EphB3 across ARMS (Rh41, **A** and Rh30, **B**) and ERMS (RD, **C**) subtypes.

### Only EphA7-Fc and ephrin-A5-Fc inhibit proliferation in hRMS cells

The divergent suites of Ephs and ephrins expressed in hRMS and the unexpected effectiveness of ephrin-A5-Fc in inhibiting hRMS proliferation raised the possibility that other Ephs or ephrins might also have an anti-proliferative effect on hRMS. We therefore tested all of the available Eph and ephrin Fc chimeras for the ability to inhibit hRMS proliferation. However, based on expression of phosphorylated histone 3 (PH3), only ephrin-A5 and EphA7 were effective at reducing hRMS cell proliferation (Figure 4).

**Figure 4.**
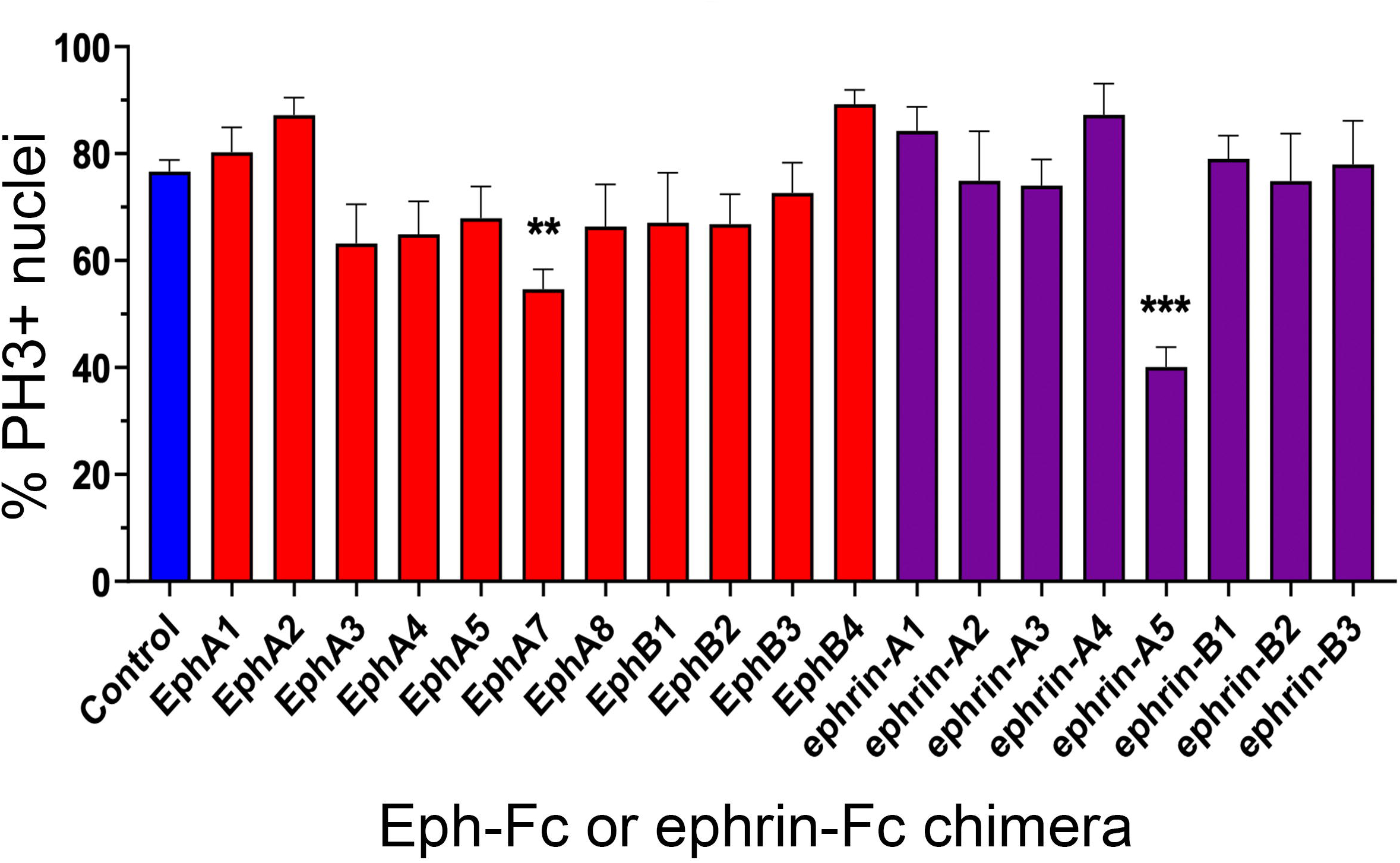
Only EphA7-Fc or ephrin-A5-Fc inhibit hRMS proliferation. Rh41 cells were cultured on all available Eph (red) or ephrin (purple) Fc chimeras for 120 hours then fixed and stained for phosphorylated histone-3 (PH3). Only EphA7-Fc (p<0.001) or ephrin-A5-Fc (p<0.0001) (consistent with data shown in Figure 1) decrease proliferation significantly.

### In hRMS, ephrin-A5 inhibits proliferation via binding to EphA8 and EphA7 inhibits proliferation by binding to ephrin-A2

Although ephrin-A5 and, to a lesser extent, EphA7 were the only Eph/ephrins chimeras able to reduce hRMS cell proliferation, the changes in localization and expression of EphA7 and ephrin- A5 in hRMS cells seemed to preclude the signaling interactions between EphA7 and ephrin-A5 we observed during developmental or regenerative myogenesis. Based on the mRNA and protein expression we had observed, we performed siRNA knockdown of 3 candidate receptors for each molecule (ephrin-A2, ephrin-A4, and ephrin-B1 were tested as potential binding parters for EphA7-Fc and EphA6, EphA8, and EphB3 were tested as potential binding partners for ephrin- A5-Fc), reasoning that knockdown of the receptor would blunt the effect of each Fc chimera on proliferation. Rh41 cells were transfected with siRNA (Silencer, Invitrogen) to each targeted molecule then treated with either EphA7-Fc or ephrin-A5 Fc. Transfected cells were scored for proliferation via PH3 staining. Of the siRNAs tested, only knockdown of EphA8 (for ephrin-A5- Fc treated cells) and ephrin-A2 (for EphA7-Fc-treated cells) abrogated the antiproliferative effects of the Fc chimera (Figure 5), indicating that these interactions are responsible for the decrease in proliferation in each case.

**Figure 5.**
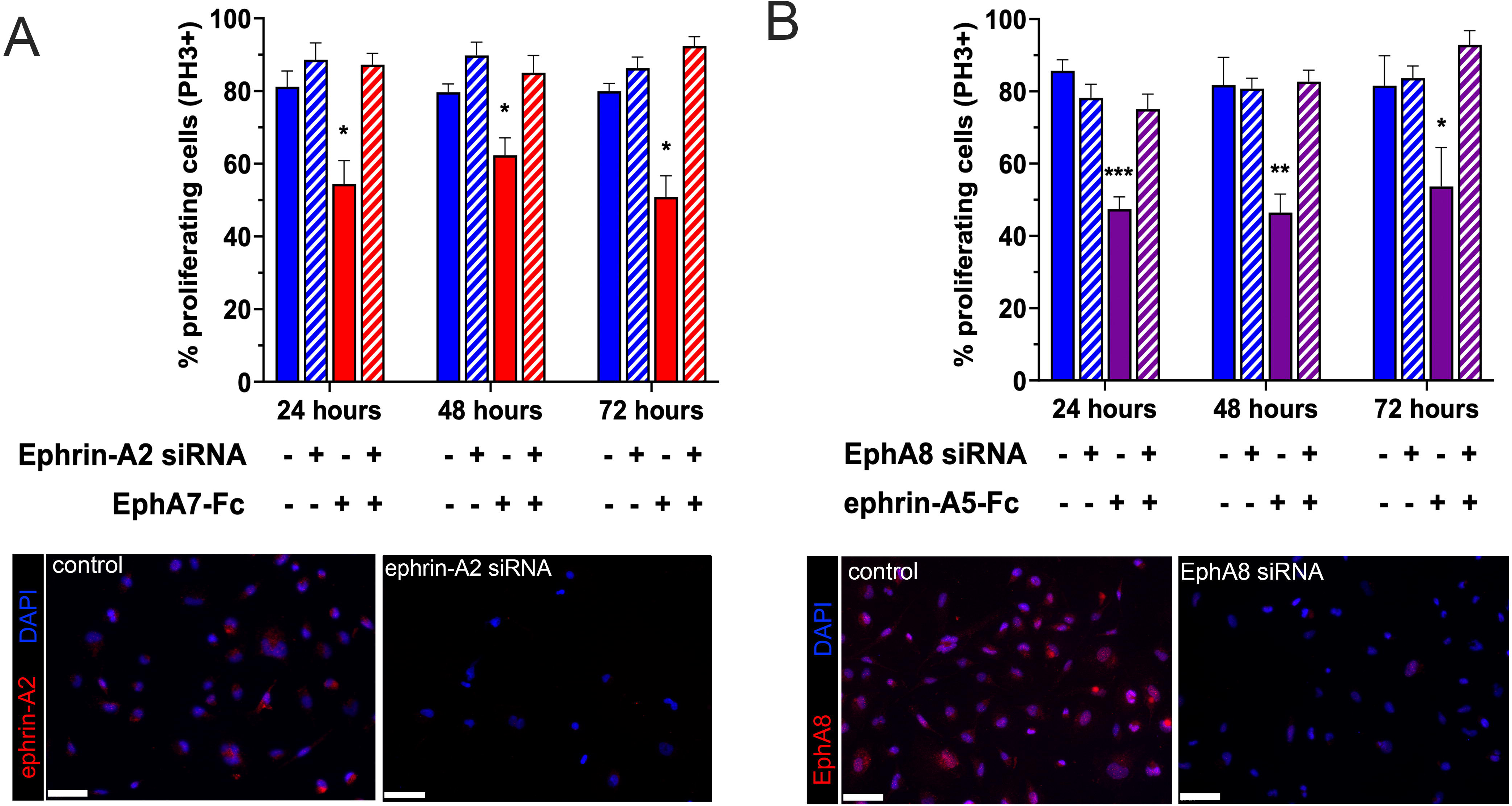
Knockdown of ephrin-A2 or EphA8 abrogates the antiproliferative effect of EphA7-Fc or ephrin-A5-Fc. siRNA knockdown of ephrin-A2 restores Rh41 proliferation to control levels even in the presence of EphA7-Fc (**A**). Similarly, knockdown of EphA8 restores proliferation even in the presence of ephrin-A5-Fc (**B**). siRNA knockdown was confirmed by immunohistochemistry for ephrin-A2 and EphA8. Bars = 50 μm; *** = p<0.0001, ** = p<0.001, * = p<0.01.

## Discussion

Rhabdomyosarcoma (RMS) is the most common solid tumor of childhood, and particularly more aggressive cancers where metastasis has occurred overall prognosis remains poor despite advances in therapeutic strategies. This study describes significantly inhibited proliferation of both ARMS and ERMS human RMS cell lines *in vitro* due to exposure to Fc chimeras of EphA7 and ephrin-A5, via different interaction partners than these molecules would bind and signal through during myogenesis *in vivo*. These data support the potential for stimulation of these pathways via soluble ephrin-A5 and/or EphA7 ectodomain as a potential differentiation therapy for hRMS.

During tumorigenesis and cancer progression, otherwise anti-proliferative molecules such as tumor suppressors and inhibitory signaling receptors are frequently inactivated, mislocalized, or repurposed to promote the tumor lifestyle [reviewed in (Wang and Li, 2014)]. We recently described aberrant localization of another Eph, EphA1, in RMS primary tumors and cell lines from three species (dog, mouse, and human) (LaCombe et al., 2022). A key finding of the current study is the aberrant localization of EphA7 to the cis-Golgi apparatus in ARMS cells, rather than the plasma membrane where it would typically exert its differentiation-promoting functions. This mislocalization likely disrupts canonical EphA7 signaling, providing a mechanistic explanation for its impaired function in RMS. The mechanism(s) by which EphA7 localization is altered in hRMS cells could be explored further, as it may provide an even more effective target for ephrin-A5 as an antiproliferative therapy.

Eph:ephrin interactions are highly promiscuous and both interactions and their downstream effects on the cell are extremely context-dependent [reviewed in (Blits-Huizinga et al., 2004; Himanen et al., 2007; Cecchini and Cornelison, 2021)]. Together with the uncertainty regarding the cell of origin for hRMS and the tendency for tumor cells to alter expression of key signaling molecules, the results indicating that EphA7:ephrinA5 signaling does not occur as it does in the context of developmental or regenerative myogenesis but that different interactions between ephrin-A5 and EphA8 and between ephrin-A2 and EphA7 have the same effect is perhaps not surprising. While such promiscuity complicates the signaling landscape, it also presents opportunities to selectively exploit these interactions for therapeutic benefit. The context-dependent nature of these interactions underscores the need for precision in targeting specific Eph:ephrin signaling axes in RMS treatment. Further research could also focus on the downstream signaling pathways that lead to decreased proliferation.

## Methods

### Cell culture

Human rhabdomyosarcoma (hRMS) cell lines RD (embryonal), Rh30 and Rh41 (alveolar) were acquired from the Children’s Oncology Group at Texas Tech University. All cell lines were grown in Iscove’s Modified Dulbecco’s Medium (IMDM - Gibco) supplemented with 20% Fetal Bovine Serum (FBS - Sigma), 4 mM L-glutamine, 1% penicillin/streptomycin, 0.1% gentamycin, and 1X Insulin-Transferrin-Selenium (ITS- Corning) at 37°C and 5% CO_2_ in a humidified incubator.

### Coverslip functionalization and cell sample preparation for immunofluorescence

Human Eph-Fc and ephrin-Fc (R&D Systems) functionalized coverslips and control laminin-coated coverslips were freshly prepared the day of seeding as previously described (Arnold et al., 2020). Briefly, HCl-washed and aminopropyltriethoxysilane (APTES, Sigma)-treated coverslips were incubated for 3 hours at 37°C with 2 mg/mL laminin (Sigma) in PBS. Laminin was then gently aspirated and coverslips were washed twice with Dulbecco’s modified phosphate buffered saline (DPBS). Half of the coverslips were then incubated for 2 hours at 37°C with a 5 µg/mL solution of EphA7-Fc in DPBS while the other half were incubated with DPBS alone. The coverslips were placed in 6-well plates with medium for 30 mins prior to cell seeding.

hRMS cells were counted using a hemacytometer and seeded at3000 cells/well. After 24, 48, 72, or 120 hours in culture cells were washed twice with DPBS and fixed with ice-cold 4% paraformaldehyde (PFA) for 10’. PFA was removed and the samples were washed with DPBS and stored in DPBS at 4°C.

### Immunocytochemistry

hRMS cells were blocked in 10% normal goat serum with 0.2% Triton X-100 for 1 hour at room temperature (RT) except for coverslips to be stained with chicken primary antibody (anti-EphA7), which were instead blocked with 10% BlokHen (Aves Labs) for 1 hour at RT. Cells were then incubated with primary antibody overnight at 4°C, washed 3X in DPBS, incubated with secondary antibody for 1 hour at RT, and washed 3X in DPBS and 1x in water before mounting with Vectashield containing DAPI (Vector Labs).

Primary antibodies from Santa Cruz Biotechnology (Dallas, TX) and used at 1 μg/mL except as noted: anti-ephrin A2: sc-912, anti-ephrin A3: sc-1012, anti-ephrin A4: sc-914, anti-ephrin B1: sc-1011, anti-Eph A5: sc-927, anti-Eph A6: sc-8172, anti-Eph A8: sc-25460, anti-Eph B1: sc- 926, anti-Eph B2: sc-28980, anti-Eph B3: sc-100299, anti-Eph B4: sc-5536, anti-PH3: sc-8656- R, 0.5 μg/mL. Primary antibodies from other sources: anti-ephrin A5, Abcam AB70114, 1:200; anti-Eph A7: made in-house, 1:300; anti-Ki67: Abcam ab16667, 1:200. Secondary antibodies were raised in goat and conjugated with Alexa fluorophores (Invitrogen) and used at 1:500.

All images were acquired and processed on an Olympus BX61 upright microscope using Slidebook software (Intelligent Imagine Innovations) or µManager software (www.micro-manager.org). Digital background subtraction was performed to remove to remove signal that was less than or equal to levels present in control samples (processed in parallel but without primary antibody) and was applied equally to the entire field.

### Image quantification and data analysis

Total number of nuclei were scored with FIJI (Schindelin et al., 2012) with the combination of commands ‘Threshold’ and ‘Analyze particles’. Stained cells were manually counted using FIJI ‘Cell counter’ tool. The number of stained cells was normalized on the number of total nuclei and converted into percentage of stained cells over total cell number in a minimum of 10 fields of view (10X magnification) per coverslip. All experiments were carried out in triplicates, except when specified otherwise.

### Statistical analysis

Where indicated, samples were compared in Graphpad Prism by ordinary two-way ANOVA, one-way ANOVA (Kruskal-Willis test) followed by Dunn’s multiple comparison test, or Mann-Whitney test.

### siRNA transfection

1,000,000 Rh41 cells were transfected with 2 μg of CMV-GFP (Clontech) and 200nM of siRNA using the Invitrogen Neon Transfection System (Voltage 1050V, width 30ms, 2 pulses). Cells were seeded in Iscove’s Modified Dulbecco’s Medium (IMDM - Gibco) complemented with 20% Fetal Bovine Serum (FBS - Sigma) and 4 mM L-glutamine. After 24 hours the cells were plated on coverslips functionalized with ephrin-A5-Fc or EphA7-Fc or on control laminin-coated coverslips. After 24, 48, or 72 hours coverslips were fixed with 4% PFA and stained for PH3 as described above.

#### RT-PCR

RNA was isolated from cell lines grown to 80% confluency on 10cm plates with the RiboPure^TM^ kit (Invitrogen) and 1/10 of the recovered RNA was used as a template for reverse transcription with SuperScript^TM^ IV RT (Invitrogen). cDNA samples were quantified by using Thermo Scientific NanoDrop One and 100ng of cDNA as template for GoTaq® Green Master Mix (Promega) 35 cycles of PCR amplification with annealing at 61°C. Intron-spanning primers used as in (Siegel et al., 2009).

## Acknowledgements

This work was supported by NIH grant AR078045 to DDWC. The authors thank Dr. Robert Arpke for statistical assistance.

---

**Supplemental Figure 1- RT-PCR screen for Eph and ephrin express in hRMS cell lines.** Intron-spanning primers were used to screen for Eph and ephrin mRNAs across hRMS cell lines. Detectable signal at appropriate molecular weights was noted for ephrin-A2, ephrin-A3, ephrin- A5, and ephrin-B1 and for EphA5, EphA6, EphA7, EphA8, EphB1, EphB2, EphB3, and EphB4 in at least one cell line.

