## Supplementary figures and images for "Ephrin-A5 or EphA7 stimulation is anti-proliferative for human rhabdomyosarcoma *in vitro*"

### Supplemental figure 1

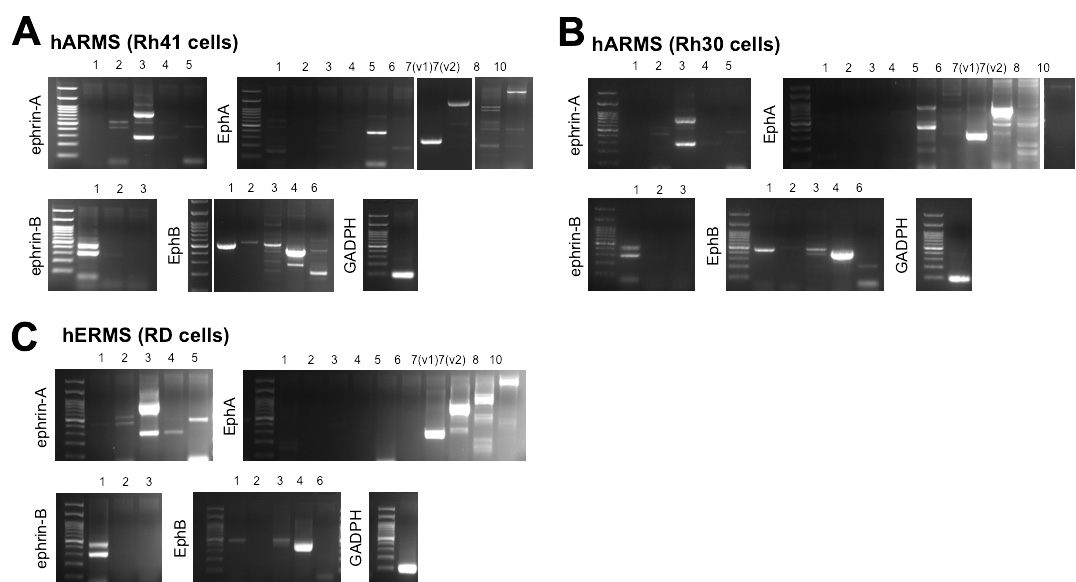
